# Testing for population decline using maximal linkage disequilibrium blocks

**DOI:** 10.1101/703595

**Authors:** Elise Kerdoncuff, Amaury Lambert, Guillaume Achaz

## Abstract

Only 6% of known species have a conservation status. Methods that assess conservation statuses are often based on individual counts and are thus too laborious to be generalized to all species. Population genomics methods that infer past variations in population size are easy to use but limited to the relatively distant past. Here we propose a population genomics approach that tests for recent population decline and may be used to assess species conservation statuses. More specifically, we study Maximal Recombination Free (MRF) blocks, that are segments of a sequence alignment inherited from a common ancestor without recombination. MRF blocks are relatively longer in small than in large populations. We use the distribution of MRF block lengths rescaled by their mean to test for recent population decline. However, because MRF blocks are difficult to detect, we also consider Maximal Linkage Disequilibrium (MLD) blocks, which are runs of single nucleotide polymorphisms compatible with a single tree. We develop a new method capable of inferring a very recent decline (e.g. with a detection power of 50% for populations which size was halved to *N*, 0.05 ×*N* generations ago) from rescaled MLD block lengths. Our framework could serve as a basis for quantitative tools to assess conservation status in a wide range of species.

## 1 Introduction

The severe and rapid changes imposed by human activities upon living organisms are suspected to be a major factor leading to short-term mass extinctions (Barnosky et al., 2011).The most comprehensive list of endangered species is the Red List of the International Union for Conservation of Nature (IUCN) (Rodrigues et al., 2006). Criteria used in the list to assess the species conservation status are based on geographical range, population trends, threats to habitat and ecology. Despite being very robust and reliable, these criteria are hard to establish for many species.To quantify the ongoing crisis for a wider range of organisms, there is a crucial need to develop quantitative measures of extinction risk to efficiently monitor species in real time and at a global scale. Previous attempts were developed to estimate quantitatively extinction rates, including by two of the present authors, based on occurrence data (Régnier et al., 2015; Ceballos et al., 2017; Sánchez-Bayo and Wyckhuys, 2019) or genetic data (from museum specimen Díez-del Molino et al. (2018); van der Valk et al. (2019)). The genetic methods measure the genetic diversity at different time to estimate the population size at these times and conclude on a general trend. The limitation of these methods is the difficulty to obtain time series data.

A handful of genomes sampled in a population at a single time point can help infer the past demography of this population (Gutenkunst et al., 2009; Li and Durbin, 2011; Excoffier et al., 2013; Harris and Nielsen, 2013; Sheehan et al., 2013; Schiffels and Durbin, 2014; Lapierre et al., 2017; Ringbauer et al., 2017; Terhorst et al., 2017; Beichman et al., 2018). In standard population genetic inferences, the periods when variations of population size can be estimated are of the order of *N*_*e*_ generations back in time. *N*_*e*_ denotes the so-called effective population size (Wright, 1931). Recent methods such as MSMC (Schiffels and Durbin, 2014) can provide inferences on more recent past but hardly scale up to large datasets of complete genomes because of their computational load. With the development of next generation sequencing, complete genomes from multiple individuals of the same species are now released routinely (Gibbs et al., 2015; Alonso-Blanco et al., 2016). Actual methods can not be applied to test for recent decline of populations, the models and methods we present in this manuscript specifically target very recent past when considering small populations and are meant to be applied to datasets of arbitrary size.

Methods using whole genome sequences to infer demography use different measures of genomic polymorphism. One of these measures is the so-called Site Frequency Spectrum, or SFS (Fu, 1995). The SFS, that is the genome wide distribution of the frequencies of polymorphic alleles in a sample of the population, is strongly distorted by the demographic history of the species (Adams and Hudson, 2004; Marth et al., 2004). SFS-based methods (e.g. Gutenkunst et al. (2009)) can handle arbitrarily large numbers of loci and genomes but disregard correlations between sites caused by genetic linkage. Using genetic linkage information may help overcoming the SFS-based methods limitations (e.g. difficulty to discriminate between different scenarios Lapierre et al. (2017) and to infer recent demography).

Recombination is the process by which two DNA sequences are intermixed to create a new sequence that combines segments of different ancestries. When two homologous regions of the genome are inherited from the same ancestor without having undergone recombination, they are said IBD: *Identical By Descent*. The probability distribution and the length of IBD regions passed through generations have been studied (Stam, 1980; Chapman and Thompson, 2003; Stefanov, 2000).

Recombination patterns are also characterized by Linkage Disequilibrium (LD). LD arises when individuals of a finite population share chunks of DNA inherited from a common ancestor (IBD blocks). Specifically, two variants located at two distinct sites are in linkage disequilibrium (LD) when their joint frequency differs from what is expected under independence. More specifically, LD is defined as the covariance 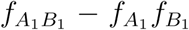, where 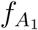 is the frequency of allele 1 at locus A (Lewontin and ichi Kojima, 1960). When 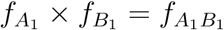, the two variants are said in complete linkage equilibrium. On average, LD decreases exponentially with genetic distance due to recombination. The pattern of LD is distorted by demography (Hill and Robertson, 1968) and thus can be used to infer the past demography of a population (Hollenbeck et al., 2016; Patin et al., 2014).

Importantly, despite the fact that breakpoints between IBD blocks are usually not observable when comparing two homologous regions, “long enough” IBD blocks can be retrieved by applying one of several recent methods to a pair of sequences (Purcell et al., 2007; Gusev et al., 2009; Browning and Browning, 2010). These methods are based on detecting long identical shared segments (Gusev et al., 2009) or shared regions that harbor multiple rare variants (Purcell et al., 2007; Browning and Browning, 2010). If two individuals share the same rare variant, they may also share the surrounding chromosomal region, particularly because rarer variants are more likely to be relatively recent. Most methods take sequencing errors into account, allowing IBD blocks to not be totally identical. The accuracy of IBD block detection depends on the algorithm used (Browning and Browning, 2013).

Some demographic inference methods are based on the distribution of lengths of inferred pairwise IBD blocks in a population. Palamara et al. (2012) have calculated the distribution of expected lengths of pairwise IBD blocks for a given parameterized demographic model. Browning and Browning (2015) have calculated the expected time to the most recent common ancestor (TMRCA) of an IBD block as a function of its length. Then they use the empirical density of IBD block lengths to estimate the distribution of TMRCA and thus the variations of effective population size through time.

Other methods use the length of identical shared segments of chromosome within a diploid individual (Hayes et al., 2003). Two identical shared segments may be inherited from a common ancestor without recombination event (and then be IBD) or may not be IBD as there are invisible recombination events that may have occurred within it. The probability that the two haplotypes of an individual share identical alleles for a given number of adjacent positions can be predicted (Hayes et al., 2003; MacLeod et al., 2009). Tools have been developed to apply these methods to infer demographic inference from genomic data (MacLeod et al., 2013; Harris and Nielsen, 2013).

Yet another approach to infer demographic history from IBD blocks is to reconstruct the genome-wide distribution of the TMRCA between two haploid genomes. In PSMC, Li and Durbin (2011) devised a Hidden Markov Model that infers the TMRCA from the positions of heterozygous sites along a pair of sequences and then estimate a step-wise demographic pattern. MSMC, the extension of PSMC (Schiffels and Durbin, 2014), analyzes the heterozygosity pattern from multiple individuals and uses first coalescence events between any two haploid genomes of the sample. These methods are computationally intensive (as of today, MSMC cannot infer the demographic history of more than 8 individuals) and pool the diversity on windows of 100 bp, that are assumed to form a single locus with two states, heterozygous or homozygous.

Importantly, the previous methods infer stepwise changes of the “effective population size” (*N*_*e*_(*t*)) that are estimated from the density of coalescence events. This motivated Mazet et al. (2015, 2016); Chikhi et al. (2018); Rodríguez et al. (2018) to propose to replace *N*_*e*_(*t*) by the more explicit *Inverse Instantaneous Coalescence Rate*. IICR only matches the instantaneous population size when the population is panmictic. It is nonetheless always possible to find a population model with constant size but spatial structure that corresponds to any IICR of a size-changing population for the TMRCA of 2 sequences (Chikhi et al., 2018). For larger samples, the joint distribution of coalescence events [*T*_2_, *T*_3_, …] can be used, in theory, to disentangle structure from demography (Grusea et al., 2019).

Existing methods for demographic inference using recombination information often use the whole genome of few individuals (less than 10) or use a smaller part of the genome. These methods only consider the joint history of two individuals (e.g. the pairwise IBD length distribution or the time of the first coalescence event between any two haploid genomes) which algorithmic complexity increases drastically with the number of individuals (e.g. detection of pairwise IBD blocks is quadratic) and generates a computational load limiting in most cases the application of the methods to a larger number of individuals. On the other hand, with few individuals, demographic inferences are unable to detect recent changes of population size.

Following the idea of Tiret and Hospital (2017), we decided to study the IBD concept extended to a multilocus segment and a larger number of individuals (*n* > 2). Some studies have been conducted on the amount of genetic material shared IBD with *n* > 2, considering closely related individuals (Donnelly, 1983; Ball and Stefanov, 2005). We extend the concept at a population level while relaxing the need for *identical* sequence (without mutation), which is why we decided to define a new term. We call ‘MRF blocks’ homologous segments that are entirely inherited from the same ancestor without recombination; these segments may or may not harbour different alleles, because of mutations. An MRF block is a segment of an alignment of haploid genomes that share the same coalescent tree. A recombination event along the sequence cuts the genome alignment into two MRF blocks, one on each side of the recombination point. By definition there is no recombination within MRF blocks so that all variants located within an MRF block are necessarily in complete linkage disequilibrium. The reciprocal is not true, as variants in complete LD do not necessarily belong to the same MRF block. MRF blocks carry the information of any recombination event that happened among the sampled individuals. As for IBD blocks, MRF blocks are usually not observable.

### Outline

We have developed a new test to detect very recent population declines of endangered species. We first consider the full length distribution of MRF blocks in a sample of haploid genomes (*n* ≥ 2). Second, as MRF blocks are not directly observable from sequence alignments, we devised a simple and efficient algorithm to chop an alignment of *n* ≥ 4 haploid genomes in Maximal Linkage Disequilibrium (MLD) blocks, that are segments which variants are in complete LD. From the length distributions of MRF blocks or MLD blocks, we devised a summary statistic to test whether a population has been declining in the very recent past. Our method is not limited by the number of genomes in the sample.

## 2 Model and Methods

In the absence of recombination, ancestral relationships between genomes can be represented in the form of a genealogical tree. Individual haploid genomes at present time are the leaves of the tree, the MRCA of these individuals is the root. The fusion of two lineages into one (a common ancestor) is named coalescence event (Kingman, 1982), hence the name of “coalescent tree”. The sum of all branch lengths that separates two genomes up to their common ancestor is the time of divergence between them, usually expressed in generations. In the Wright-Fisher model with constant population size *N*, branch lengths measured in number of generations scale like *N*. In particular, if we define *T* as the total length of the coalescent tree, the expectation of *T* is proportional to *N*. Large populations generate coalescent trees with deep nodes, whereas small populations have shallow coalescent trees.

In the presence of recombination, two loci of an alignment have the same coalescent tree only if no recombination event happened since their MRCA. We name MRF block, a maximal interval along the alignment of sites sharing the same coalescent tree. MRF blocks are consequently separated by recombination points, corresponding to recombination events. It is standard to assume that conditional on the total length *T* of the coalescent tree of a site, the length *L* of its MRF block is exponentially distributed with rate *ρT*, where *ρ* is the recombination rate (expressed in a arbitrary unit proportional to Morgan). Then for a fixed *ρ, T* and *L* are negatively correlated: recombination is more likely to occur in deep trees, which thus are carried by shorter blocks. As mentioned above, as *T* is proportional to *N*, MRF blocks are also shorter in larger populations. More accurately, because the law of *T/N* does not depend on *N*, neither does the law of *NL* (for *n* = 2 it alludes to results of Carmi et al. (2014)). In other words, if population 1 has size *N*_1_ and population 2 has size *N*_2_, the distribution of MRF block lengths in population 2 can be deduced from that in population 1 by a scaling factor *N*_1_*/N*_2_, both populations having identical demography otherwise. For example if *N*_2_ = 2*N*_1_, the MRF blocks in population 2 are twice smaller than those of population 1.

Note that for a given *N* the lengths (*L*_1_, *L*_2_, …) of successive adjacent blocks have the same distribution, but they are not independent, because the coalescent trees of adjacent MRF blocks are not. The dependencies between these trees is encoded in the so-called Ancestral Recombination Graph (ARG) (Griffiths and Marjoram, 1997). Because these dependencies have a complex structure (Wiuf and Hein, 1999), a popular way of approximating them is the Sequentially Markovian Coalescent (SMC) (McVean and Cardin, 2005; Marjoram and Wall, 2006). This approximation neglects coalescences between lineages with no overlapping ancestral material and assumes Markovian dependencies of coalescent trees along the sequence: the genealogy of an MRF block only depends on the genealogy of the adjacent ones.

Although genealogies of different MRF blocks are not independent, they are asymptotically independent as the distance between them increases.

Throughout this article, we use msprime to generate MRF blocks directly from the ARG (Kelleher et al., 2016) but very similar results were obtained with a local SMC implementation. We assume constant recombination and mutation rate along the genome. We simulated the alignment of *n* = 10 haploid genomes at present time.

### Demographic scenario

We consider a single change of population size (Fig 1a). Here *N*_*t*_ represents the population size at time *t, t* = 0 is the present time and positive values represent the past. We denote by *κ* the ratio of the two sizes: *κ* = *N*_∞_*/N*_0_, and by *τ* the time at which the population size changes in coalescent units of *N*_0_ generations. If *κ* = 1, *N*_∞_ = *N*_0_: there is no change. If *κ* = 10, *N*_∞_ = 10*N*_0_: the population size has been divided by 10, *τ N*_0_ generations in the past.

**Figure 1:**
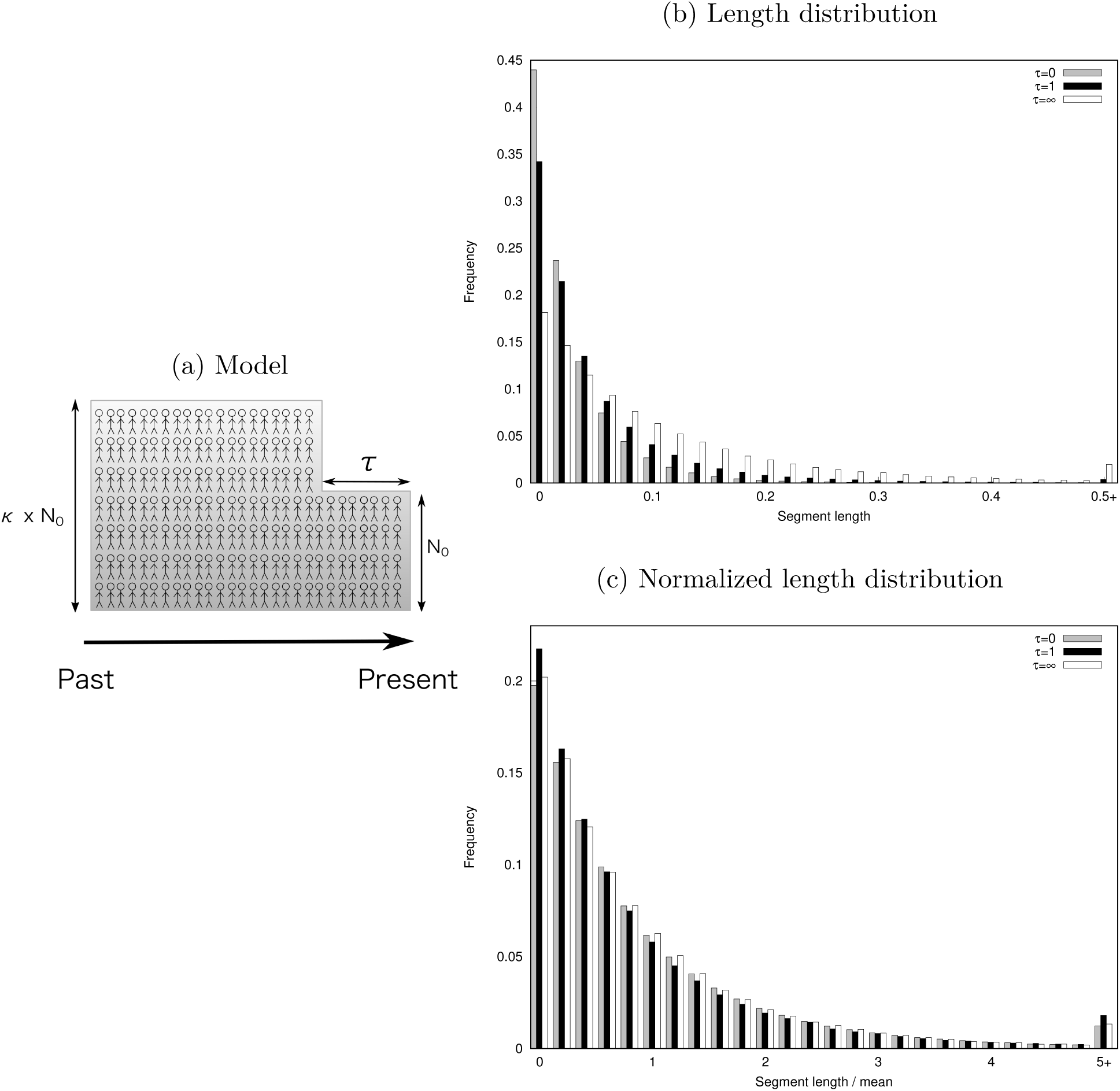
Impact of the demography on the distribution of MRF block lengths. (a) The demography considered here is a sudden size change tuned by 3 parameters : *N*_0_, the actual population size, *τ* the date of decline (in backward time) and *κ* the strength of decline. Time is expressed in *N*_0_ generations. (b) Distribution of *L* for *ρ* = 1, *κ* = 3 with *τ* = {0, 1, ∞}. When *τ* = 0 (grey) or *τ* = ∞ (white), population size is constant. (c) Distribution of *L*′ = *L/E*[*L*] under the same values of *ρ* = 1, *κ* and *τ*. In case of a decline, the distribution is overdispersed, with an excess of both short and long normalized MRF blocks.

## 3 MRF blocks

### Distribution of block lengths

#### Impact of population decline on tree length

For declining populations (*κ* > 1), the coalescent trees have two distinct time scales: a first one for the shallow part of the tree (*t* < *τ*), expressed in *N*_0_ generations, and a second one for the deep part of the tree (*t* > *τ*) that is expressed in *κN*_0_ generations. When the declining population tree is compared to a standard coalescent tree (constant population size), it has shorter external branches if the reference time scale is expressed in *κN*_0_ generations *or* longer internal ones if the reference time scale is expressed in *N*_0_ generations. When it is compared to a reference tree with population size chosen so as to have the same *T*_*MRCA*_, its external branches are too short *and* its internal branches are too long. Similarly, the distribution of the total length *T* of the tree is overdispersed when compared to the length of the standard coalescent tree with the same mean.

#### Impact of population decline on lengths of MRF blocks

For a declining population, the distribution of the length *L* of MRF blocks will depend not only on *ρ* and *N*_0_ but also on *κ* and *τ*. As the tree relative branch lengths are distorted and the distribution of *T* is overdispersed, so is the distribution of *L*. In a declining population, the distribution of *L* can be seen as a mixture of the two distributions that correspond to the two population sizes, *N*_0_ and *κN*_0_. The strength of the decline (*κ*) tunes the difference between the distributions; the date of decline (*τ*) tunes in what proportion the two distributions are mixed. When *τ* → 0 (practically, *τ* < 10^−4^ times *N*_0_ generations for a sample size *n* ∈ [10, 100]), the distribution of *L* is indistinguishable from that of block lengths in a population with constant size equal to *κN*_0_. At the opposite, for *τ* → ∞ (practically, *τ* > 10 times *N*_0_ generations for a sample size *n* ∈ [10, 100]), the distribution of *L* is indistinguishable from that of block lengths in a population with constant size equal to *N*_0_. As a result for *τ* ∈ [10^−4^, 10], the distribution of *L* has an excess of MRF blocks smaller than the *N*_0_ reference and an excess of MRF blocks longer than the *N*_∞_ reference (Fig 1b). The small blocks correspond to the trees which total length *T* is mostly driven by the distant *N*_∞_ time scale and the long ones to the trees which total length *T* is mostly driven by the recent *N*_0_ time scale.

As mentioned in the previous section, in a population with constant size *N*, the distribution of *L*, briefly denoted *L*_*N*_, scales like 1*/N*, in the sense that the distribution of 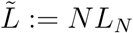 does not depend on *N*. In particular, the distribution of *L*′ := *L/E*[*L*] does not depend on *N* in a population with constant size and follows the law of 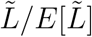. However, the distribution of *L*′ is distorted when there is a size change. For a declining population, the distribution of *L*′ is overdispersed, it has an excess of small blocks (*i.e.* less than 0.2) and an excess of long blocks (*i.e.* more than 5), as can be seen on Fig 1c.

Note that we always have *E*[*L*′] = 1, but here *E*[*L*] has

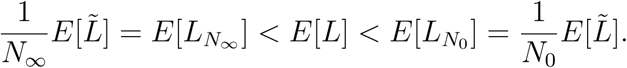

As the block distribution of *L*_*N*_ is a mixture of the one of 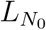 and 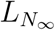, *E*[*L*] is bounded by 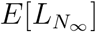 and 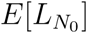 that depend on the population size.

## 4 MLD blocks

### 4.1 Definition

All recombination events are not directly visible in a genome alignment. First, adjacent MRF blocks may have coalescent trees sharing the same topology and the same branch lengths, so that mutations occurring on either tree show exactly the same pattern on either block. Second, adjacent MRF blocks may have coalescent trees sharing the same topology but not the same branch lengths, so that mutations occurring on either tree display the same bipartitions (compare the second and third tree in Figure 2). Third, even if two adjacent MRF blocks have trees with different toplogy, it is possible that branches distinguishing these topologies do not carry mutations (see the second block in Figure 2).

Importantly, recombination events that happen between the two oldest lineages do not impact the topology of the tree, so are never detectable because they do not impact the possible bipartitions.

**Figure 2:**
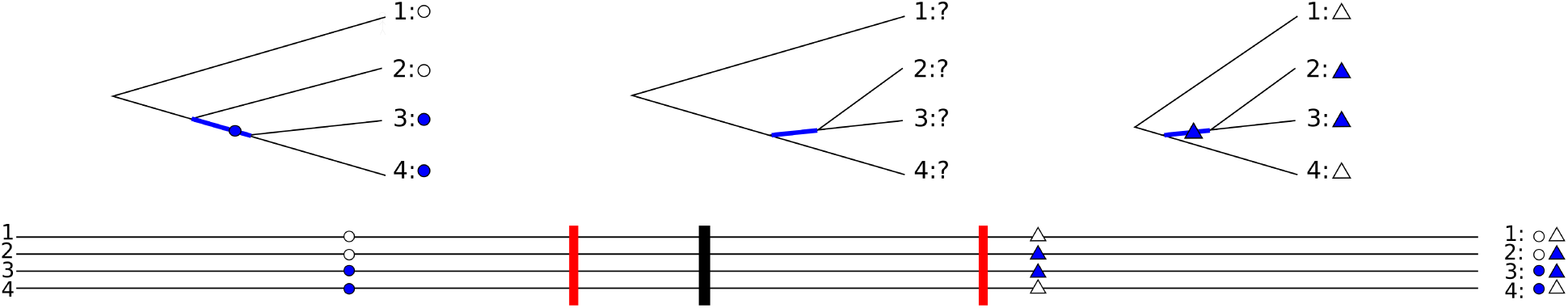
Detection of recombination in three MRF blocks. The four lines represent four haploid genomes, circles and triangles are mutation events, red lines are the true recombination events delimiting MRF blocks and above each MRF block is represented its true tree. Mutation events occur on certain lineages as represented on the trees. The first recombination event generates an incompatibility between the blue branches of the first two MRF blocks, but as no mutation occurs on the second MRF block, this recombination event cannot be detected. The second recombination event does not change the topology of the tree and thus this second event cannot be detected either. However, the first and the third MRF blocks carry mutations that are not compatible; thus a minimum of one recombination event can be inferred between the two mutations, as indicated by a vertical thick black line arbitrarily placed in the middle.

A possibility used in the literature to detect breakpoints between MRF blocks is to detect the changes in the density of polymorphic sites along the sequence due to the change of coalescent tree (like in PSMC, Li and Durbin (2011)).

Here we used instead the incompatibilities between bipartitions displayed by polymorphic sites to place the minimal number of recombination events on the alignment. Two bipartitions are said incompatible when they are not compatible with a common tree.

In what follows, we will assume that the mutation rate *µ* is constant through time and along the genome.

#### 4.1.1 The four-gamete test

From now on, we assume that each site can be hit at most once by a mutation, so that a polymorphic site is always bi-allelic, an assumption known as the “infinitely-many sites model”. The four-gamete test (Hudson and Kaplan, 1985) serves to detect incompatibilities between bipartitions displayed by two polymorphic sites. For any two biallelic sites (A/a and B/b) there are at most four gamete haplotypes in the population (A-B A-b a-B and a-b). Under the infinitely-many sites model, the four possible haplotypes cannot be observed in a sample if the two sites share the same genealogy. Then if the four possible haplotypes are observed in the sample, a recombination event must have occurred between them – but not necessarily the other way round. This property can be used to compute a lower bound for the number of recombination events in a genome alignment (Hudson and Kaplan, 1985) or even to estimate the recombination rate (Hey and Wakeley, 1997). We used it to compute and place the minimal number of breakpoints in a genome alignment.

Two polymorphic sites are said *incompatible* if the four possible haplotypes are present in the sample. When a sequence of adjacent polymorphic sites contains no pairwise incompatibility, we speak of a sequence of compatible sites. Note that a sequence of compatible sites are in complete linkage disequilibrium. We thus define an MLD block, for *Maximal Linkage Disequilibrium* block, as any maximal sequence of compatible sites.

We now explain how to extend this notion originally designed for haploid genomes (or phased diploid genomes) to an unphased diploid genome, that is, a diploid sequence lacking the linkage information. For an unphased diploid genome, the two original haplotypes can be determined if the diploid genome is homozygous at at least one the two sites:

- When the genome is homozygous at both loci (A/A-B/B), both haplotypes must be A-B.
- When the genome is homozygous at one locus and heterozygous at the second one (A/A-B/b), the haplotypes must be A-B and A-b.

The four-gamete test can then be extended to a sample of unphased diploid genomes by saying that two sites are incompatible in this sample if they are incompatible in the subsample of haplotypes that have been inferred thanks to the previous remark. When the haplotype is ambiguous, the sites are considered compatible and do not bring more information about a recombination event.

#### 4.1.2 The chopping algorithm

We used the four-gamete test to detect incompatibilities in the genome alignment and to chop it into MLD blocks (Fig 3). To avoid computing the full matrix of pairwise incompatibilities between all polymorphic sites of the genome, we only compute the incompatibilities for sequences of *P* adjacent polymorphic sites (by default *P* = 150). Each pair of incompatible sites (*i, j*) defines an interval that contains at least one MLD breakpoint. To place the MLD breakpoint, we seek the shortest interval that is sufficient to explain the incompatibilities.

**Figure 3:**
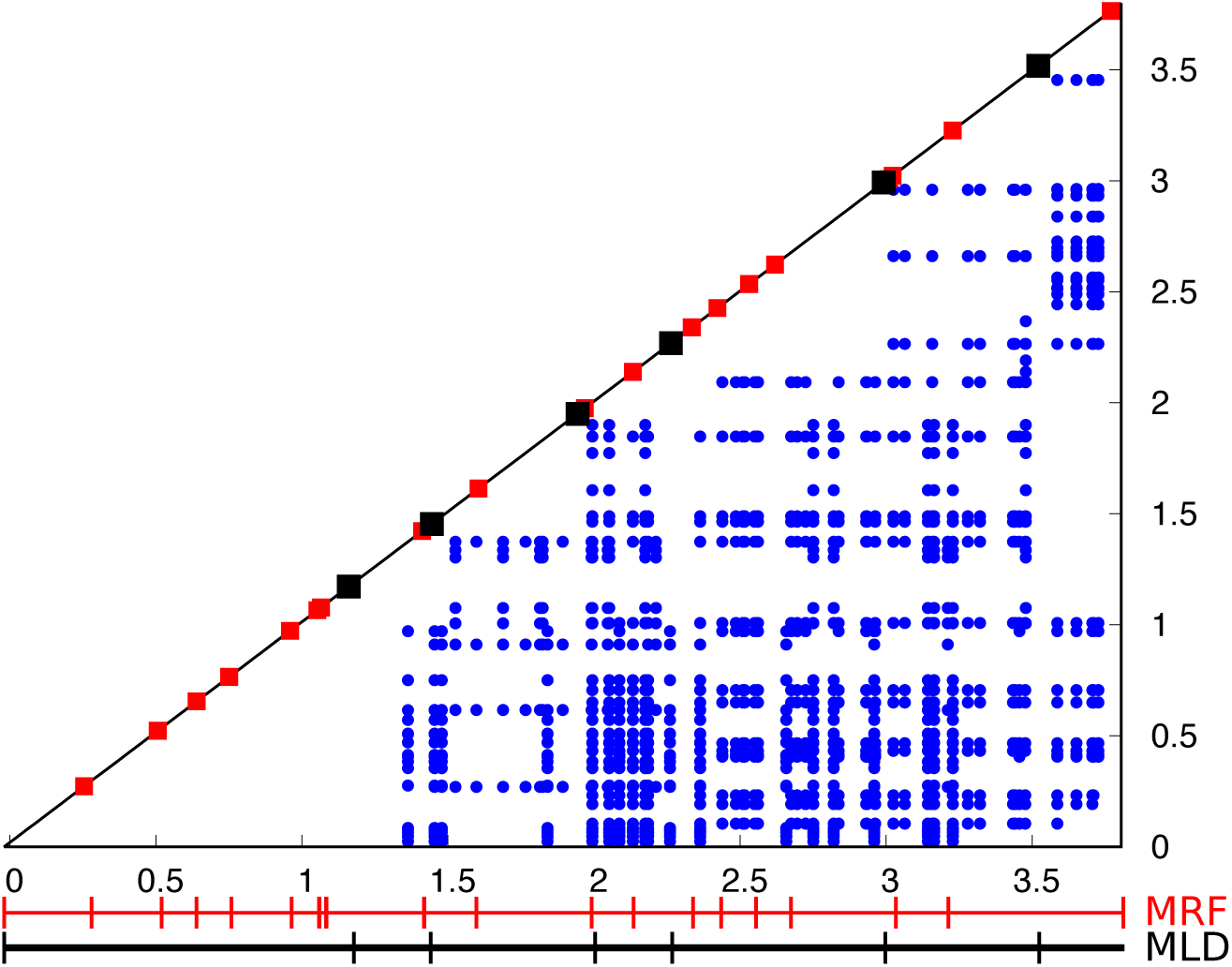
The incompatibility matrix and the chopping algorithm. X and Y-axis are positions on the genome alignment. Blue dots represent a pair (x,y) of incompatible sites. The red squares are the true positions of recombination events (MRF breakpoints) and the black squares are MLD breakpoints inferred by the chopping algorithm.

##### Algorithm

We retrieve all intervals and sort them in increasing order of site positions along the genome (first by *i* the first site position and when equal, by *j* the second site position). As we scan two times the list of intervals, the algorithm complexity is linear with the number of polymorphic sites:

1. **Discarding and shortening.** For this step, we scan the list in reverse order, from the last (*N*) to the first interval. (The algorithm can be done in the forward order, the distribution will be slightly different but it will not affect the study.) Each interval containing another entire interval is discarded: for two intervals (*i*_*N*_, *j*_*N*_) and (*i*_*N*−1_, *j*_*N*−1_), if *i*_*N*_ ≤ *i*_*N*−1_ ≤ *j*_*N*−1_ ≤ *j*_*N*_, then (*i*_*N*_, *j*_*N*_) is discarded. When two intervals overlap, they are replaced by their intersection (the two original ones are discarded): for the two intervals (*i*_*N*_, *j*_*N*_) and (*i*_*N*−1_, *j*_*N*−1_), if *i*_*N*−1_ ≤ *i*_*N*_ ≤ *j*_*N*−1_ ≤ *j*_*N*_, both are replaced by a new interval (*i*_*N*_, *j*_*N*−1_), that is then compared to (*i*_*N*−2_, *j*_*N*−2_)…
2. **Positioning.** From the final list of disjoint intervals, we place an MLD breakpoint at the middle of each interval.

MLD breakpoints partition the genome alignment into MLD blocks.

### 4.2 Length distribution

The distribution of the length *L*_*c*_ of a typical MLD block does not only depend on the distribution of *L* (MRF block length) but also on the fraction *p* of recombination events that are detected. This fraction increases with the ratio *µ/ρ*, as illustrated in Figure 4a. When many mutations occur in two different MRF blocks (*µ* ≫ *ρ*), the probability that they occur on incompatible branches of their respective coalescent trees increases and so does the detection efficiency, up to a point of saturation due to cases when these MRF blocks share the same tree topology. The number of sampled individuals also impacts the efficiency of detection (Fig 4a): the larger the sample size, the higher the probability to observe incompatible mutations. The four-gamete test for unphased diploid genomes has obviously less power to detect recombination than for phased genomes (Fig 4a).

**Figure 4:**
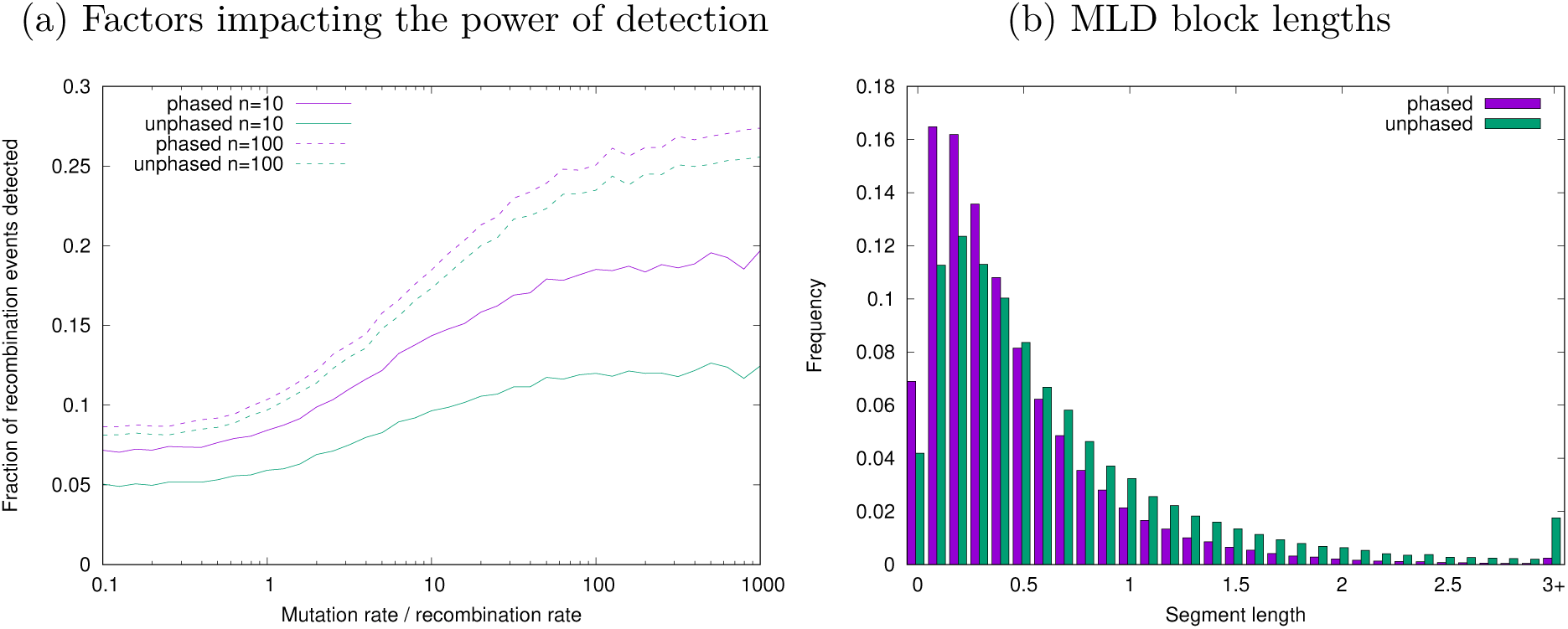
Detection of recombination events and its impact on MLD block length (*L*_*c*_) distribution under a constant population. (a) Fraction of recombination events that are detected as a function of *µ/ρ* for different sample sizes (*n* = 100, dashed lines and *n* = 10 plain lines) and for phased (purple) or unphased (green) diploid genomes. (b) Distribution of MLD block lengths for phased (purple) and unphased (green) diploid genomes in a population of constant size (*µ* = 10, *ρ* = 1, *n* = 10).

The lower the power to detect recombination, the longer the MLD blocks. In particular, phased genomes have smaller MLD blocks than unphased ones (Fig 4b). Furthermore increasing the sample size results in more detectable recombination points and thus smaller MLD blocks. In Figure 4b, the average block length, in our arbitrary unit for *n* = 10 phased haploid genomes is 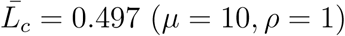. Considering smaller sample size will result in larger MLD blocks (e.g. 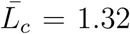 for *n* = 5). This implies that the total number of blocks can be limiting for small sample size, and that these long blocks will be harder to detect in scaffolds of partial genomes. In (very) large samples, MLD blocks are shorter: 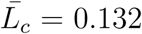 for *n* = 600 (ten times smaller than for *n* = 5) and 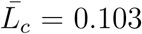 for *n* = 6, 000. On a side note, the theoretical pitfall of having too small “undetectable” blocks can always be overcome by subsampling.

Here, we consider the block lengths normalized by the average length 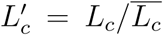. Similarly to the MRF blocks, the distribution of 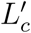 does not depend on the value of *N* but does depend on the demographic scenario (Fig 5). However, it still depends on our ability to detect recombination and so on the ratio *µ/ρ* and *n* the number of sampled individuals. To compare distributions, it is then important that they have the same ratio *µ/ρ* and the same *n*.

**Figure 5:**
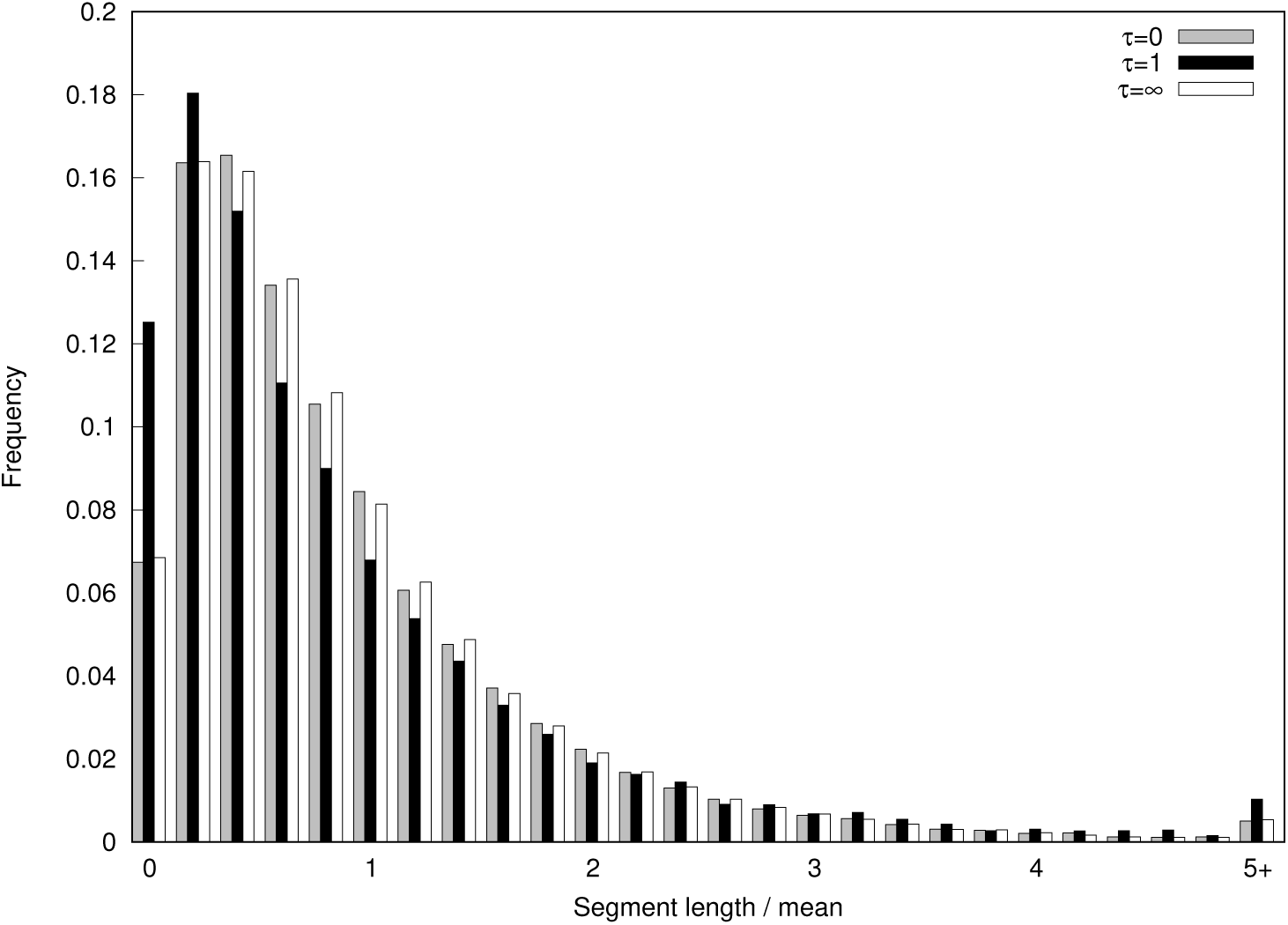
Distribution of 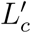 for a population of constant size (white, *N* = *N*_0_ = 1 and grey, *N* = *κN*_0_=3) and for a declining population (black for *τ* = 1) with *ρ* = 1, *µ* = 10 and *n* = 10.

Similar to what we have observed for MRF blocks, a declining population exhibits both an excess of small blocks 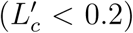 and large blocks 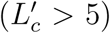 (Fig 5). The shape of the distribution of 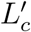 (Fig 5) differs from the one for *L*′ (Fig 1c): MLD blocks are longer than MRF blocks. Indeed, they contain a variable number of MRF blocks and below a certain size, MRF blocks are not detectable as recombination points at the edges of an MRF block can be detected only when mutations have occurred inside the block. MLD blocks are always longer than MRF blocks.

## 5 Statistical tests for population decline

### 5.1 Test

To test for population decline, we use the excess of small and large blocks that we observe when comparing samples from a declining *vs* a constant population size. More specifically, we compute the fraction of blocks which normalized length is either smaller than 0.2 or larger than 5, both in the case of MRF blocks (*f* = *f*_*L′<0.2*_ + *f*_*L*′ >5_) and of MLD blocks 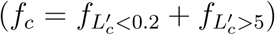. To set an empirical threshold value under *H*_0_, we simulate 10,000 genomes of 10^5^ MRF blocks under a constant population size for 10 haploid genomes and compute both 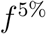 and 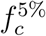 as upper limits for one-tailed tests: *f*^5%^ = 0.214236 and 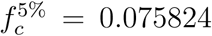. As the threshold is empirical, simulations need to be redone for a change in sampled size or in null model. Time needed for simulations depends on the algorithm/software used and the specific features of the model.

### 5.2 Power

To assess the power of this test, we simulated 1,000 replicates under population decline (*H*_1_ with various *τ* and *κ*) and report the fraction of runs where 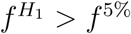 for MRF blocks or 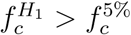 for MLD blocks. When the power is 1, the decline was significant in all runs. When the power is 5%, the decline is not detectable, the test can not differentiate *H*_0_ and *H*_1_.

Without surprise, results show that the power of the test to detect population decline depends on both the decline strength (*κ*) and the date of decline (*τ*) (Fig 6). For both tests based either on MRF or on MLD blocks, the power outreaches the 5% risk only for a range of *τ*. The type I error of the test is 5% as expected. For both tests, the range of detection is wider when the decline is stronger (compare dashed to solid lines in Fig 6). The surprise is that the test based on MLD blocks (*f*_*c*_) detects more recent declines than the test based on MRF blocks (*f*). Therefore, we recommend using the *f*_*c*_ test when searching for very recent decline even if MRF blocks are known (which is generally not the case).

**Figure 6:**
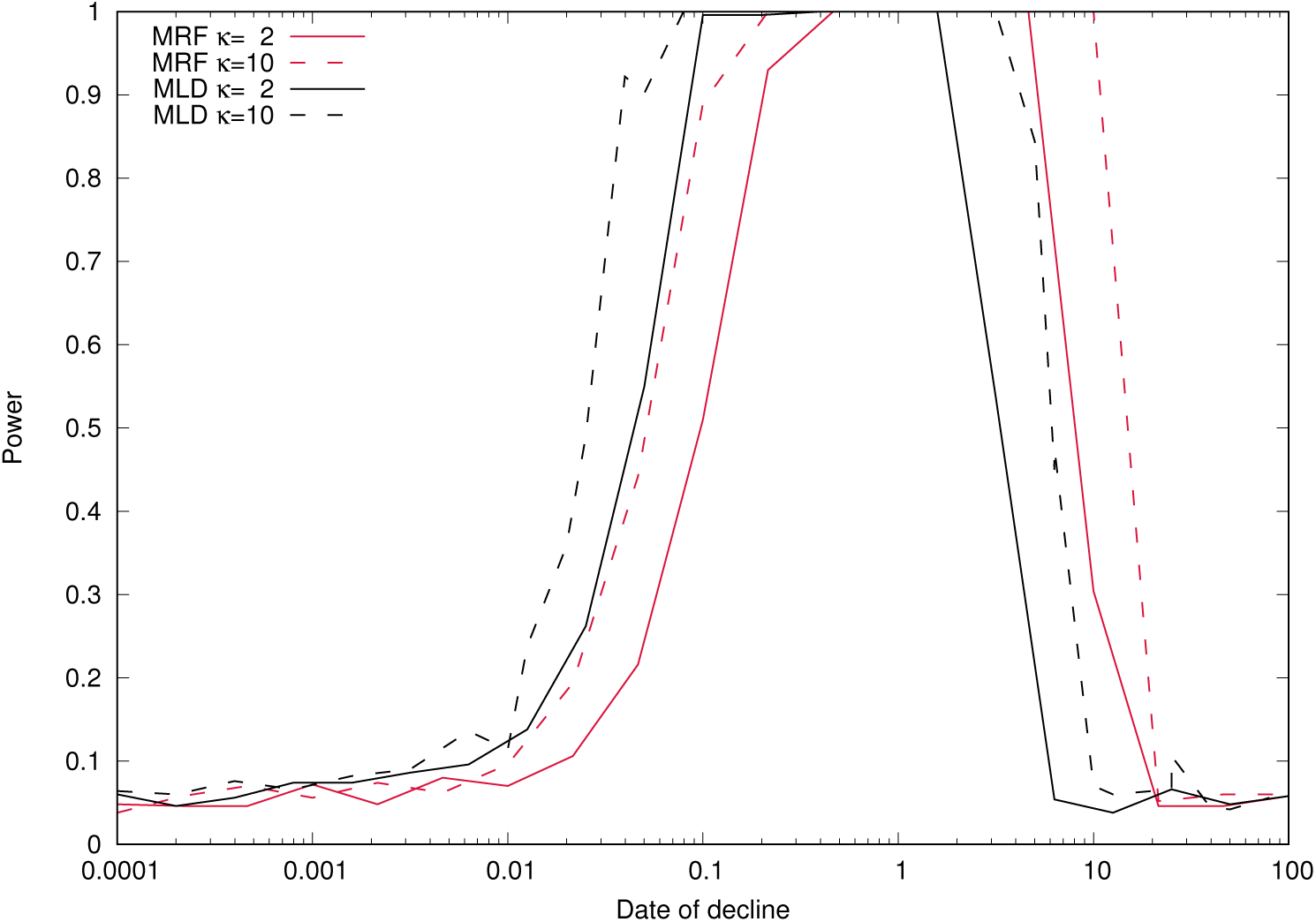
Power to detect population decline. The test based on MRF blocks (*f*) is pictured in red, whereas the one based on MLD blocks (*f*_*c*_) is represented in black. We assess the power of the two tests for *κ* = 2 (plain line) and *κ* = 10 (dashed line) with *τ* ∈ [0.0001, 100].

## 6 Application to data: the case of the western lowland Gorillas

### 6.1 Handling the low quality of real genomes

Genomic data sets often include sequencing errors and regions that are not genotyped. Consequently, the *f*_*c*_ test cannot be run as is on these data sets. We present some modifications to our test to handle the poor quality of data. We show in this section that adjustments can be made to get the information from the 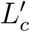 distribution.

#### Simulations with lower quality

Difficulties in applying the *f*_*c*_ test to real data sets can stem from the low quality of DNA sequences. We replicated in the simulated genomes the two main issues, namely the interruptions of DNA tracts and the absence of genotyping for some SNPs in some individuals.

DNA tract interruptions truncate MLD blocks and make their detection difficult. The number, size and location of these interruptions will have an effect on the detection of MLD blocks and thus will alter the 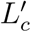 distribution. To handle the effect of interruptions, we placed the interruptions at the same positions in our simulated chromosome as in the real chromosome.

As for the partial genotyping issue, we artificially lowered the genotyping quality in the simulated chromosomes. We used the empirical distribution of missing individuals (e.g. chr1 of *Gorilla gorilla*, Fig 7) to pick random positions in the simulated chromosome and erase the genotypes of some individuals.

**Figure 7:**
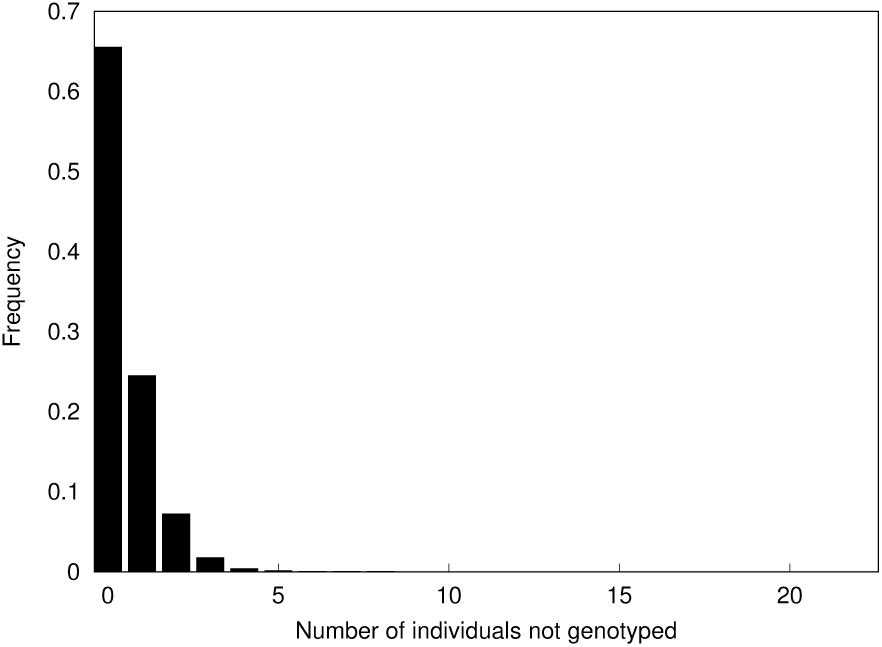
Distribution of the number of individuals not genotyped per SNP on chromosome 1 of the Gorillas dataset (Prado-Martinez et al., 2013).

#### Mutation rate and recombination rate

To cope properly with the issue of genotyping, we simulated chromosomes with the same number of mutations and the same MLD length mean as in the studied data set. We use the Watterson estimator (Watterson, 1975) for the mutation rate and fixed the recombination rate so that simulated and real chromosomes had the same average length of MLD block.

### 6.2 Application to Chr1 of *Gorilla gorilla gorilla*

We applied this methodology on chromosome 1 of twenty-three unrelated western lowland Gorillas (*Gorilla gorilla gorilla*) from the Great Ape Genome Project (Prado-Martinez et al., 2013). The chromosomes have 247,249,719 base pairs. The 23.1% of sites that are considered “low coverage”(Prado-Martinez et al., 2013) divide the chromosome alignment into 6,277,293 uninterrupted stretches. The 5,388,083 interruptions due to a single site were not considered as *interruptions*. To speed up simulations, we considered stretches longer than 499 sites, as smaller stretches often carry no entire MLD block. We chopped chromosome 1 using a window of 150 polymorphic sites, into 7,082 MLD blocks with an average length of 307.897 bp.

#### Distribution of 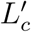

The distribution of 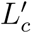 for our sample of gorilla sequences has an excess of small and long MLD blocks compared to the 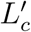 distribution of a constant population with the same characteristics (same number of mutations and same average length of MLD blocks) (Fig 8). The excess of small blocks is even larger than what we see in simulated declines (see above). The truncation of long MLD blocks due to the inclusion of low quality of genotyping can potentially inflate this excess.

**Figure 8:**
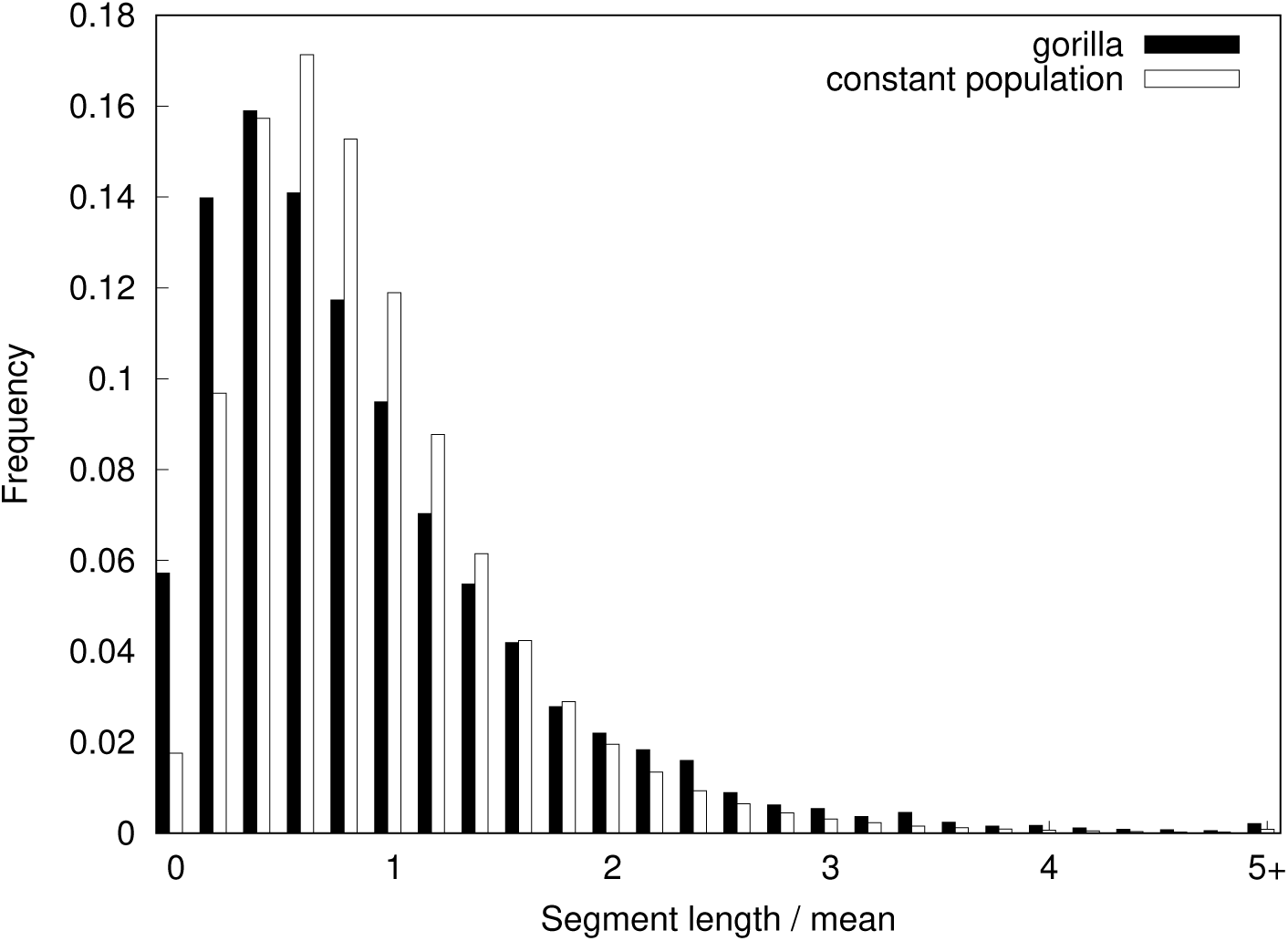
Distribution of 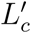 for a population of constant size (white, mutation rate= 0.000375, recombination rate= 0.012) and for the chromosome 1 of the gorillas (black)

With the low quality of genotyping and the chosen mutation and recombination rates, the threshold value 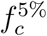 is 0.041627. As we measure 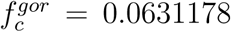 for the gorillas, the test significantly rejects *H*_0_. However, it is possible that other designs of similar tests (tweaking the lower or upper bounds) may be more relevant to analyze demography from low quality chromosome alignments.

However, and this may be even more important, misspecifications of the model can also make the test significant. Among all, we have chosen to explore the impact of recovery after the decline and of spatial structure.

## 7 Misspecification of *H*_1_

To appreciate how the *f*_*c*_ test, that was specifically designed to detect population decline, is sensitive to other violations of *H*_0_, we explored their sensitivity to a scenario of bottleneck (decline followed by recovery) and to a scenario with structure but no demography.

### 7.1 Bottleneck

In the bottleneck scenario, we model a population that experienced a sudden strong decline (*κ* = 10) at time *τ* = 1 in the past and recovered to its original size after a duration of *x* ∈ [0, 1]. If *x* = 0, there is no population decline. If *x* = 1, the population has not recovered and the bottleneck scenario is identical to our original *H*_1_. When the bottleneck lasts long enough (*x* > 0.02), it is detected by the *f*_*c*_ test (Fig 9b). On the contrary, when the bottleneck is too short (*x* < 0.02), the distribution of 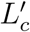 is similar to the one under *H*_0_ (Fig 9a). This shows that even if the population has recovered, the signal of decline will be observable in the excess of short and long MLD blocks.

**Figure 9:**
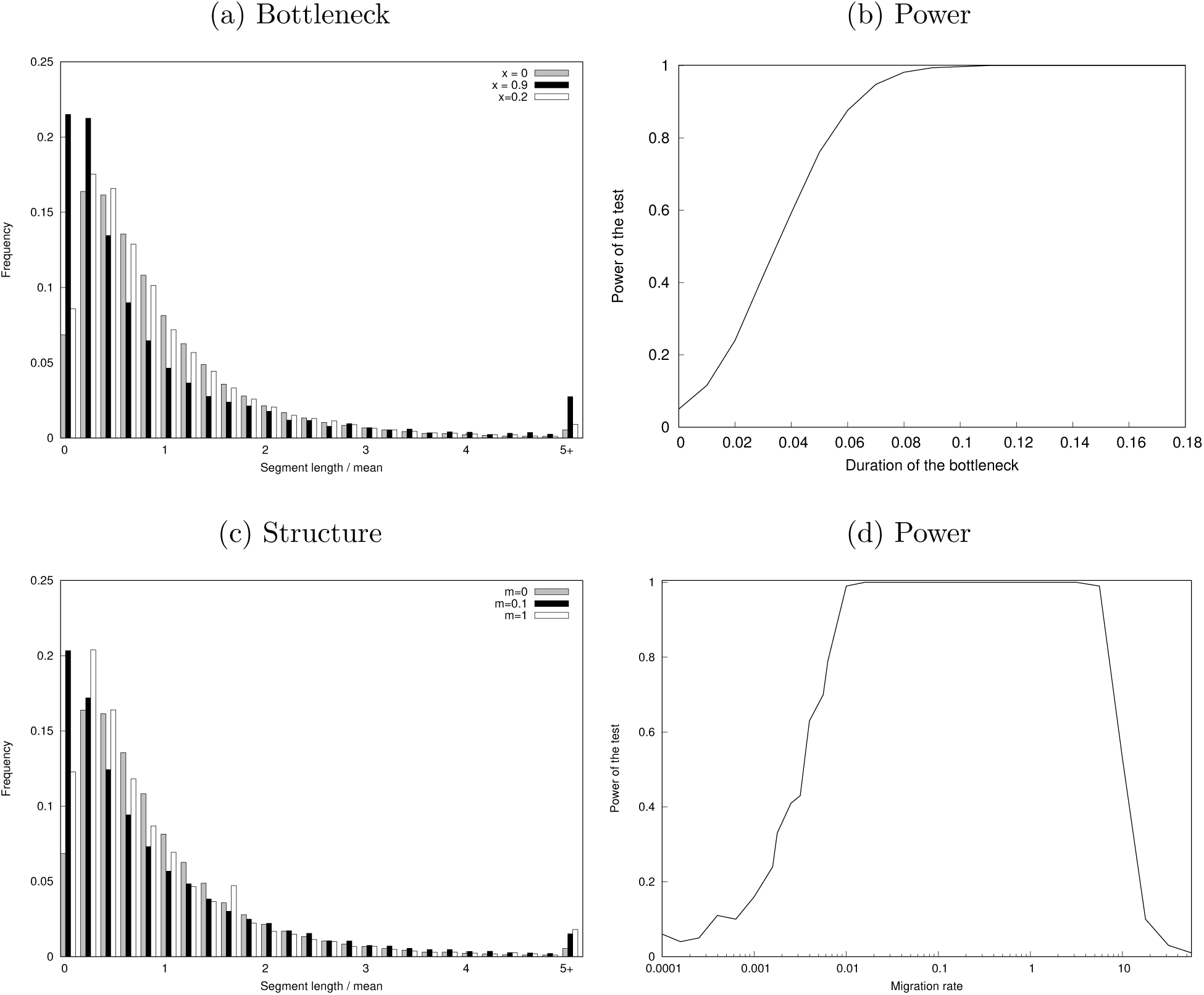
Distribution of 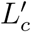 and power of the *f*_*c*_ test under two alternative scenarios. (a) In the bottleneck scenario, the population size is constant equal to *N* except during the period [1 − *x*, 1] (measured in units of *N* generations backwards from the present) during which it equals *N/*10, with *x* = {0, 0.2, 0.9} (*x* = 1 corresponds to decline without recovery). (b) Power of the *f*_*c*_ test on the bottleneck population as a function of the duration of the bottleneck, *x* ∈ [0, 0.18]. (c) In the island-mainland scenario, the population size is constant equal to *N* (island) and receives migrants at individual rate *m* = {0, 0.1, 1} from a population of size 10*N* (mainland). (d) Power of *f*_*c*_ test as a function of *m*, the migration rate from the mainland to the island.

### 7.2 Island-mainland structure

Structured populations generate signals of population size change, even when the population is stationary (Mazet et al., 2015). For example if the size of the sample is *n* = 2, for any population model with spatial structure, there exists a model without structure but specifically designed variations of population size which has the same distributions of coalescence times (Mazet et al., 2016). We consider here a larger sample size (*n* = 10, as in the other scenarios). We assume that genomes are sampled from an island with population size *N* and the island receives migrants with individual rate *m* from the mainland, which has population size 10*N*. The shape of the distribution of 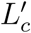 is impacted by the migration rate and the ratio of population sizes between the island and the mainland (data not shown). When the migration rate gets too small (*m* < 0.001) or too large (*m* > 10), the distribution of 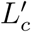 is the same as under *H*_0_ (Fig 9c). For intermediate values (*i.e.* 0.001 < *m* < 10), an a excess of short and long blocks will be observed (Fig 9d). However, the shape of the distribution of 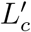 is visually different from the one of a declining population, that is, the excess of small blocks is higher than under a declining scenario. For a value of *m* = 0.1, the proportion of blocks between 0 and 0.1 times the mean is significantly higher than the proportion of blocks between 0.1 and 0.2 times the mean, which is not observed for the distribution of block lengths under decline. This suggests that the distribution could be used to differentiate between the effects of demography and structure.

## 8 Discussion

We have explored the impact of demography, more specifically recent population decline, on the pattern of recombination in a sample of *n* genomes, where *n* ≫ 2. We have shown that the distribution of the distances between recombination breakpoints (MRF block lengths) is strongly affected by the demography. More specifically, a decline will result in an overdispersion of the distribution, that is, a relative excess of short and long blocks. As most recombination breakpoints are difficult, and sometimes impossible, to detect in a sequence alignment, we have proposed to restrict ourselves to the ones that can be detected using the four-gamete test. These detectable breakpoints delineate blocks in full linkage disequilibrium that we named MLD blocks.

Although different from the distribution of MRF blocks, the distribution of MLD block lengths is also overdispersed when the population has been declining recently. Using simple tests based on an excess of small and long blocks (*f* and *f*_*c*_), one can detect declines for a wide range of different dates and strengths.

Surprisingly, the *f*_*c*_ test based on MLD blocks has more power for very recent declines (*τ* ≈ 0.01) than the *f* test based on MRF blocks. The past demography of the population impacts the distribution of the length *L* of MRF blocks but also the fraction *p* of MRF breakpoints that also correspond to MLD breakpoints. When recombination occurs at distant times when only *k* ≪ *n* ancestor lineages are present (*i.e.* the most ancient times of the tree), it rarely produces incompatibilities detectable with the four-gamete test (never when *k* = 2). For a declining population, these ancient lineages have longer branches than the ones of a constant population scenario, so that recombination events occur more frequently in these lineages. This results in a smaller *p* for declining populations and thus in more numerous (ancient, small) MRF blocks per MLD block. The relative abundance of long recent MLD blocks becomes thus more important in the distribution. This effect fades away for distant declines. In summary, the effect of recent declines on the *L*_*c*_ distribution is the result of both a change in the *L* distribution and a change in the fraction *p* of breakpoints detected, which can explain the difference in power between the *f* and the *f*_*c*_ tests.

We also show that using the *f*_*c*_ statistic, the decline can still be detected even if the population has recently recovered its original size (bottleneck scenario). Finally, we showed that local sampling of a small deme with constant size also leads to rejection of *H*_0_ for *f*_*c*_ but that the distribution of block lengths seems distorted in a way that can help distinguish the two scenarios. We leave this for future work.

One interesting advantage of using the *f* and *f*_*c*_ tests are their efficiency in computing time, such that they can scale up to a very large sample of long genomes. For example, the chopping of the entire human chromosome 1 (1,636,975 SNPs) for a sample of 10 unphased genomes takes 16 seconds on a laptop (with an Intel Core i7 processor running macOS High Sierra). This very short computing time is an interesting asset of this test compared to other methods based on variations (e.g. in SNP density) induced by recombination events (Li and Durbin, 2011; Palamara et al., 2012; MacLeod et al., 2013; Harris and Nielsen, 2013; Browning and Browning, 2015). The main choice that influences the computation time of the chopping algorithm is the number of sites considered for the chopping window. An increase in the chopping window size will increase the number of sites to test for incompatibility.

In the theoretical assessment of the *f*_*c*_ test, we have made the assumption that the recombination rate is constant along the genome and that entire genomes are aligned. Let us discuss the limits of these assumptions.

First, the recombination rate is known to vary along the genome, especially in regions of high recombination known as recombination hot-spots. It could be possible to integrate these variations via the knowledge of the recombination map. Indeed, if the recombination rate is twice higher in a given region of the genome, MRF blocks will be twice smaller, so we can correct this distortion by multiplying all MRF block lengths by 2.

Another issue of the test based on MLD block lengths is the need of whole genome data. For normalisation of the MLD block distribution, the average length of a block is needed. If the whole distribution of MLD block is not available, it can compromise the estimate of the average length, and so can compromise the test based on the normalised distribution. The test requires genome data with good SNP quality for all the individuals.

The *f*_*c*_ test is a genome-wide approach that can detect population decline that started even very recently, down to orders of *τ* = 0.01*N*_0_, where *N*_0_ is the current (effective) population size. This corresponds to very recent times, in particular when considering endangered populations. For example, there are approximately 600 mature mountain Gorilla individuals alive (IUCN Red List of 31 July 2018). Assuming that the current effective population size is a third of the mature individuals, *N*_0_ ≈ 200, the *f*_*c*_ test will detect decline as recent as 0.01 ∗ 200 = 2 generations ago. Great apes populations (Bonobos, Chimpanzees, Orangutans) have been sequenced (Prado-Martinez et al., 2013) and are actively re-sequenced (Gordon et al., 2016). The coverage used to sequence the data currently available is not high enough to apply our test. To infer MLD blocks, the sequenced DNA tracts need to be uninterrupted. Using these whole-genome data in higher quality, we will be able to confirm their decline thanks to the *f*_*c*_ test.

Giant pandas have a ‘vulnerable’ conservation status in the Red List. Recently they have seen their population increased (around 500 mature individuals, from IUCN website 2019). Applying the *f*_*c*_ test on some genomes of theirs (Zhao et al., 2013) sequenced in higher quality, will give some precise information on their demography. As the test is influenced by duration and strength of a bottleneck, the strength and the date of the increase in the population size impact the result of the test. Applying the *f*_*c*_ test to mammals with approximately known demography will be interesting to verify the method. However, the real asset of this test is its possible application to a much wider range of organisms. Whole-genome data start to become more and more common for non-model, non-vertebrate organisms like honeybee (Wallberg et al., 2014), as well as organisms with no conservation status such as mimicry butterflies (Zhang et al., 2017).

The chopping algorithm detects incompatibilities among trees along the genome. All the sites in a MLD block are compatible with one topology. We developed the algorithm to detect a recent change in population size. However, its use is not limited to population demography. Conflicting genealogies are also present in phylogenetic inference (Maddison, 1997). This algorithm could be used to partition the genome according to compatible trees before estimating the trees.

Recombination and mutation events leave a joint imprint on genomes which depends notably on the demography of the population. Their frequency and locations carry information about the past history of this population (decline, bottleneck, structure…). Using MLD breakpoints to chop genomes gives insights into this history and may be used to gain further information on other aspects impacting the frequency of recombination events through time and along the genome (e.g. hitch-hiking due to selection).

## Acknowledgements

E.K. is funded by the PhD program ‘Interfaces pour le Vivant’ of Sorbonne Université. G.A., A.L. and E.K. thank the *Center for Interdisciplinary Research in Biology* and the *Fondation François Sommer* for funding.

